# Closed-loop beta stimulation enhances beta activity and motor behaviour

**DOI:** 10.1101/2025.10.07.680961

**Authors:** Min Wu, Zeyu Xu, Melanie K Fleming, Nic Shackle, Lara Biller, Faye Tabone, Pei-Ling Wong, Caroline Nettekoven, Andrew Sharott, Catharina Zich, Charlotte J Stagg

## Abstract

Movement-related beta event-related synchronization (ERS) has been linked to motor control and learning, showing potential as a therapeutic target for those with movement deficits, such as stroke survivors. However, whether directly modulating beta ERS can causally influence motor performance remains unclear, largely due to the lack of methods designed to specifically target this neural activity. To address this gap, we developed a novel behaviourally-driven, closed-loop transcranial alternating current stimulation (tACS) approach to target movement-related beta ERS during a visuomotor adaptation task. We found that the behaviourally-driven, closed-loop beta-tACS specifically enhances beta ERS without affecting beta event-related desynchronization (ERD). Critically, this targeted enhancement significantly improves retention of motor adaptation. These findings establish a causal relationship between beta ERS and motor behaviour and highlight the potential of behaviourally-driven beta-tACS as a therapeutic approach for improving motor function in clinical populations characterized by impaired beta activity.

## Introduction

From performing key-hole surgery to playing Tchaikovsky’s Piano Concertos, humans are uniquely able to learn and perform skilled movements. Even in our daily lives being able to type, dress, and feed ourselves requires precise timing of neural activity to learn and refine motor patterns. The extent of motor impairment in patients with neurological disorders has been linked to the degree of disruption in this neural activity, making it a highly promising target for therapeutic rehabilitation. Here, we present the first evidence that targeting the neural activity underpinning motor patterns using closed-loop non-invasive brain stimulation can modulate that neural activity, leading to behavioural improvements and offering a promising approach for therapeutic interventions in clinical populations.

Human movement is dependent on changes in neural activity within the cortico-basal ganglia circuits in the beta frequency range (13-30 Hz). Specifically, movements are characterised by a decrease in movement-related beta activity during movement execution (event-related desynchronization, ERD) and an increase following movement termination (event-related synchronization, ERS) ^1,2^. The amplitude of movement-related beta activity patterns relates to movement kinematics and motor learning in animals and in healthy adults ^3–6^, and alterations of movement-related beta activity are observed in a range of neurological conditions. For example, the reduction of the beta ERD mediates the disordered movement characteristic of Parkinson’s Disease, and treatments that alleviate motor symptoms appear to do so via restoring movement-related beta activity ^7–9^. Similarly, the beta ERS is decreased after stroke and increases during motor recovery, with people who recover motor function having a stronger beta ERS than those who do not recover motor function ^10–12^. Indeed, non-invasive brain stimulation approaches aiming at improving motor recovery after stroke, also increase the beta ERS ^13,14^. However, the causality of these relationships has been impossible to determine: does improved movement lead to increased ERS, or does increased ERS lead to improved movement?

We address this question here by developing a novel, behaviourally-driven closed-loop non-invasive brain stimulation approach that triggers transcranial alternating current stimulation (tACS) in real time to target a specific brain state. tACS applies sinusoidally-varying electrical current through the scalp, and can modulate endogenous neural activity at specific frequencies ^15^. Traditionally tACS involves applying continuous stimulation irrespective of the ongoing brain state ^16–18^. However, recent studies indicate that the effectiveness of brain stimulation depends on the underlying brain state at the moment of stimulation ^19,20^, among other factors such as participant anatomy ^21^. This has driven the development of closed-loop, state dependent stimulation that adjusts stimulation in real time. We have developed a behaviourally-driven closed-loop beta-tACS approach, where stimulation was triggered by the participant’s behaviour. Behaviourally-driven closed-loop brain stimulation may offer advantages over neurally-driven approaches by targeting behaviourally-related neural activity more directly, which can potentially lead to more effective outcomes. In our case we use the end of the movement, in line with the beta ERS (Fig. 1), to trigger beta-tACS aiming at enhancing the beta ERS and thereby improve motor control.

**Figure 1.**
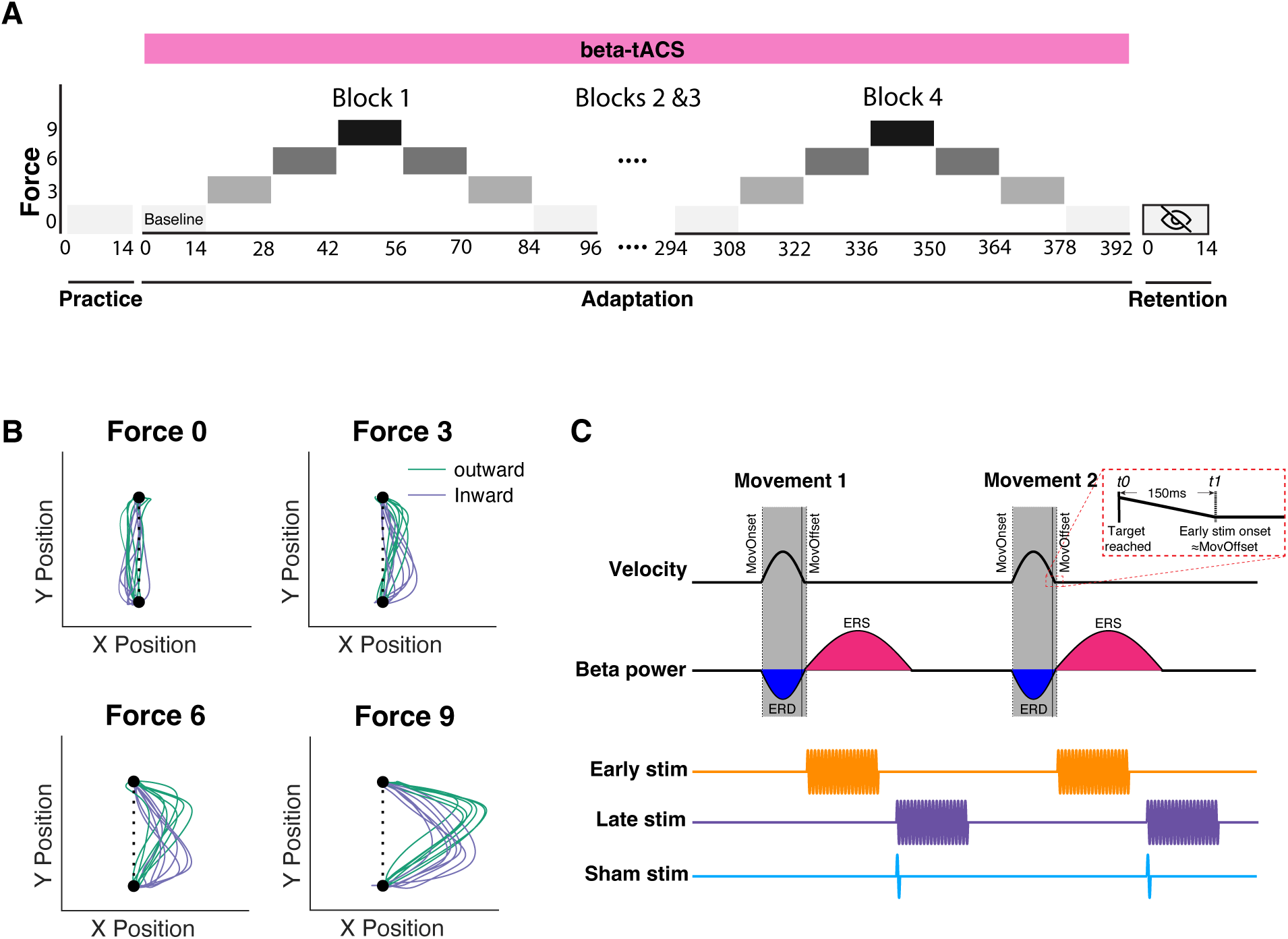
Experimental setup and stimulation protocol. **(A)**. Schematic of target-reaching motor task. The task consisted of three phases: a practice phase, an adaptation phase with a lateral force field perturbation, and a retention phase with no force field and without visual feedback. **(B).** Example movement trajectories from a single participant at different force field magnitudes (0, 3, 6, and 9). Green trajectories depict outward/upward movements and purple trajectories depict inward/downward movements. **(C).** Overview of the behaviourally-driven closed-loop tACS. In the early stimulation condition (orange), 20 Hz tACS was triggered following movement offset (i.e., *t_1_,* 150 ms after target reached [*t_0_*]) and lasted 2 s. In the late stimulation condition (purple), tACS was triggered 2.5 s after movement offset and lasted 2 s. In the sham condition (blue), stimulation was triggered 2.5 s after movement offset but lasted only 100 ms. Time *t₀* was defined as the moment that the cursor was movement into the target area and time *t₁* was defined as the moment that the cursor remained continuously within the target area for 150 ms.

Here, 24 young healthy adults completed three separate sessions in a double-blind, sham-controlled crossover design involving three stimulation conditions: (i) an early stimulation condition, delivering 2s of beta-tACS at movement offset; (ii) a late stimulation condition, delivering the same stimulation 2.5 s after movement offset; and (iii) a sham stimulation condition, delivering brief (100 ms stimulation) also at 2.5 s after movement offset. To investigate the stimulation effects on both behavioural and neural levels, we used a target-reaching visuomotor adaptation task with a Kinarm robotic system to assess motor behaviour, and simultaneously recorded electroencephalography (EEG) to measure beta activity in stimulation free periods. We hypothesized that behaviourally-triggered, closed-loop beta tACS delivered at movement offset (i.e., the early stimulation condition) would enhance beta ERS in the sensorimotor cortex and facilitate the acquisition of a motor skill.

## Results

### Behaviourally-triggered beta tACS enhances post-movement beta ERS but not ERD

We first wanted to investigate the effects of behaviourally-triggered beta tACS on the post-movement beta ERS. EEG was recorded during the performance of the target-reaching visuomotor adaptation task and was analysed from those trials where tACS was not applied. As expected, after participants reached the target, there was a robust increase in beta-band power (beta ERS; 15-30 Hz) over the left sensorimotor area, contralateral to the upper limb performing the movement (Fig. 2).

**Figure 2.**
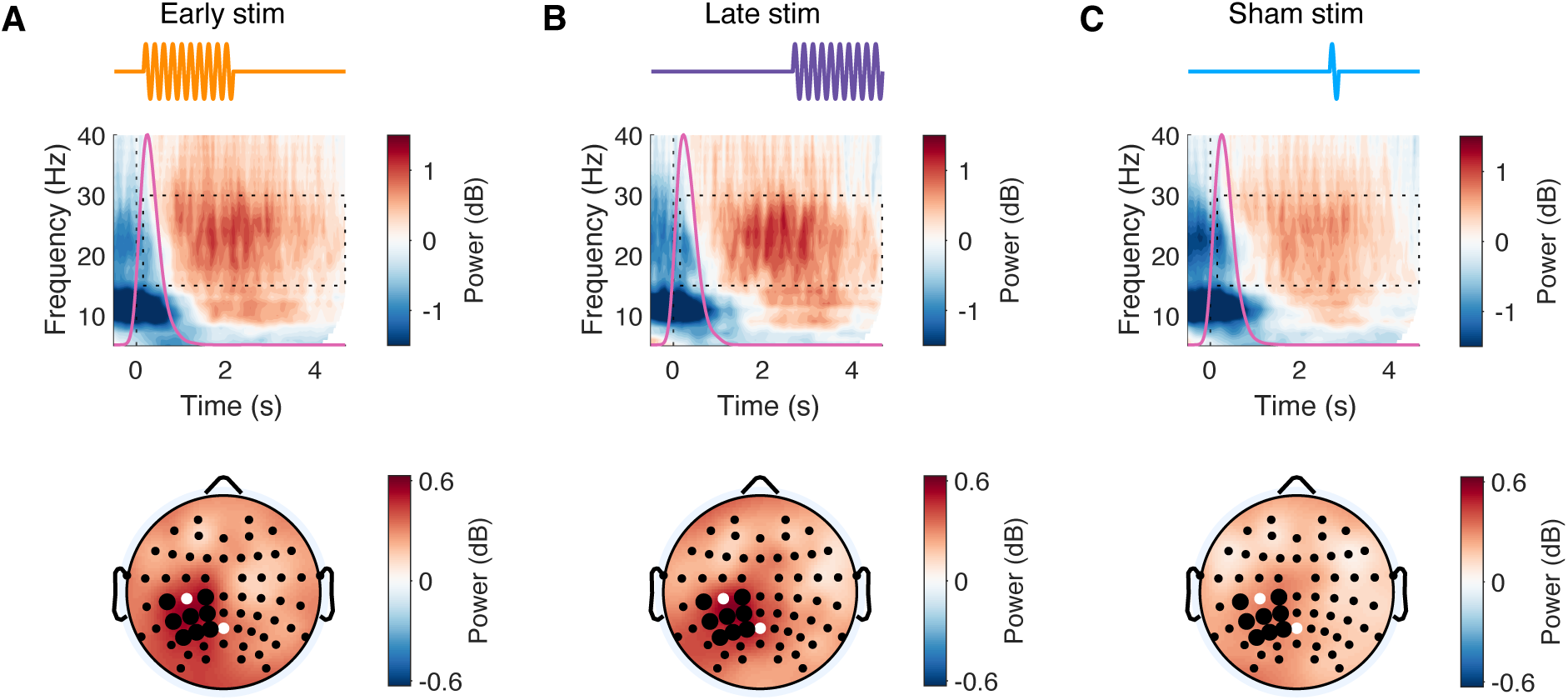
Post-movement time–frequency responses across stimulation conditions. **(A)**. TACS and time-frequency response aligned to target reached (*t₀* = 0 s) for the early stimulation condition. The power was averaged across sensors covering the contralateral sensorimotor cortex (highlighted in bold in bottom panel). The magenta curve shows the distribution of the movement offsets across participants. The black dashed rectangle indicates the time–frequency window used to quantify post-movement beta ERS. The scalp topography shows a contralateral distribution of beta ERS. Black dots mark electrodes in the predefined sensorimotor area; white dots indicate stimulation electrodes (C3 and Pz). **(B).** Similar as (A), for the late stimulation condition. **(C).** Similar as (A), for the sham stimulation condition. All EEG analyses were performed on non-stimulated trials, and trials containing stimulation artefacts were rejected.

To determine whether stimulation modulated motor-related beta activity, we performed cluster-based permutation testing on the time-frequency responses in the contralateral sensorimotor cortex (Fig. 3A), and on the mean beta power in the contralateral sensorimotor cortex (Fig. 3B). These analyses demonstrated that beta activity in the early stimulation session differed significantly from beta activity in the sham session primarily within the early post-movement window, whereas beta activity in the late stimulation session differed significantly from the sham session in the late window.

**Figure 3.**
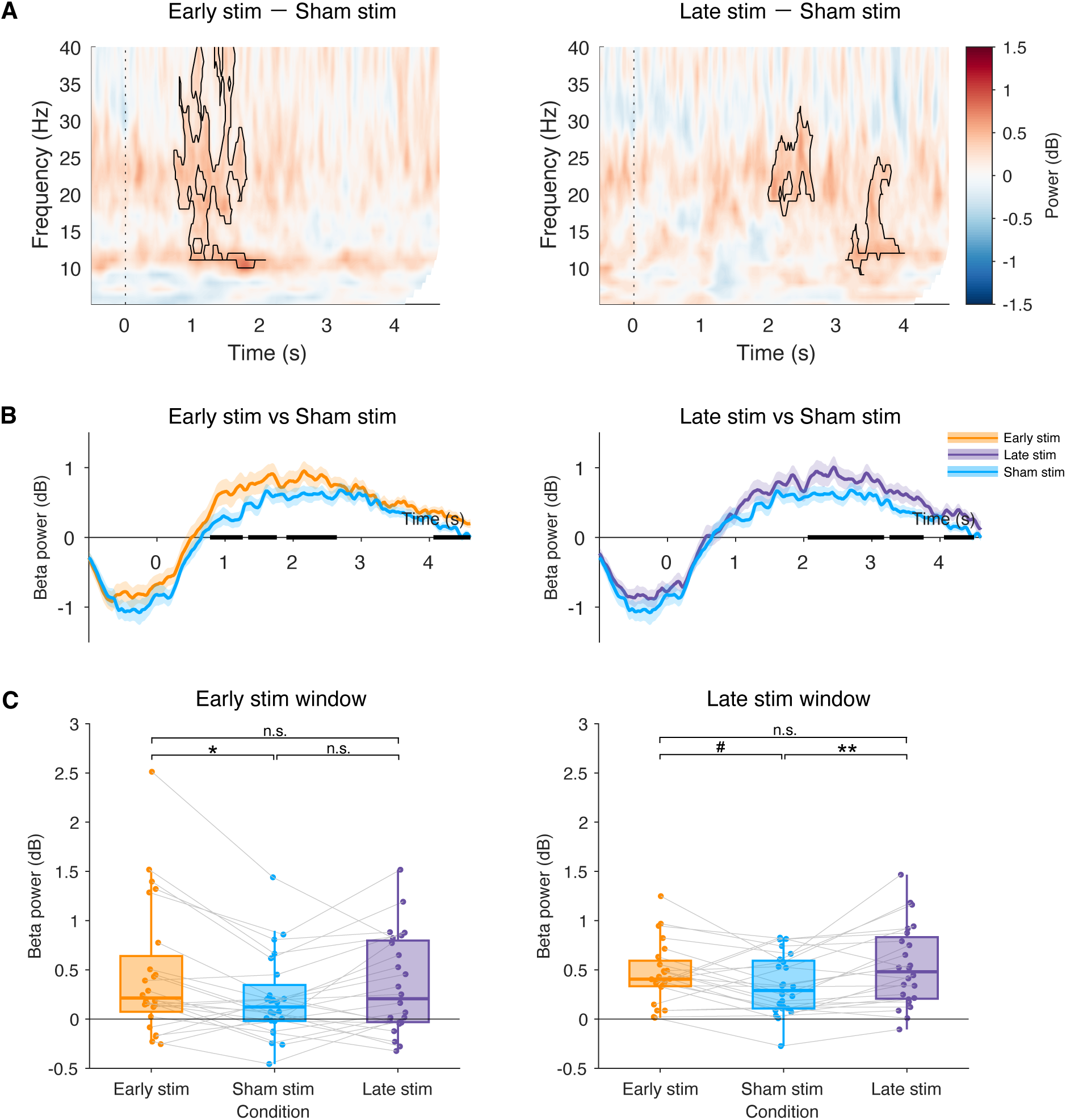
Modulation of post-movement beta activity by stimulation conditions. **(A)**. Time frequency response difference in the sensorimotor cortex between conditions. The black outlines indicate significant clusters with p < 0.05 by cluster-based permutation test. **(B).** Time course of beta power (15–30 Hz) averaged over the contralateral sensorimotor area, aligned to target reached (*t₀* = 0 s). Left: comparison between early (orange) and sham stimulation (blue). Right: comparison between late (purple) and sham (blue) stimulation. Shaded areas indicate s.e.m. Black horizontal bars indicate time intervals with significant differences between conditions, determined by cluster-based permutation testing (*p* < 0.05). **(C).** Mean beta power extracted from the early (left) and late (right) stimulation windows. Each dot represents an individual participant; grey lines link participant data across conditions. Box plots show the median and interquartile range (IQR). Compared to sham stimulation, beta ERS was significantly enhanced following early stimulation in the early window and following late stimulation in the late window. n.s. non-significant, ^#^ *p* < 0.1, * *p* < 0.05, ** *p* < 0.01.

These findings were further supported when we calculated mean beta power within two stimulation-aligned time windows: the early stimulation window (0–2 s relative to *t₁* [∼movement offset]) and a late stimulation window (2.5–4.5 s relative to *t₁*), and compared across stimulation conditions. In the early stimulation window, beta ERS differed significantly across stimulation conditions (χ²(2) = 7.874, *p* = 0.019; Fig. 3C, left). Beta ERS was significantly higher in the early stimulation session compared to sham (*t*(24) = 2.734, *p* = 0.026, FDR-corrected), with no significant differences observed between late stimulation and sham (*t*(24) = 1.598, *p* = 0.175, FDR-corrected), or between early and late stimulation (*t*(24) = 1.136, *p* = 0.261, FDR-corrected).

In the late stimulation window, there was also a main effect of stimulation condition (χ²(2) = 10.61, *p* = 0.005; Fig. 3C, right). Beta ERS was significantly enhanced in the late stimulation session compared to sham (*t*(24) = 3.168, *p* = 0.008, FDR-corrected), though no significant difference was observed between the early and late stimulation sessions (*t*(24) = –1.270, *p* = 0.210, FDR-corrected).

To determine whether these stimulation-related changes were specific to the beta ERS, we investigated the amplitude of the beta ERD in a similar way. As would be expected, there was a significant decrease in beta-band power (i.e., beta ERD) during movement, distributed bilaterally over sensorimotor area (Extended Data Fig. 1). However, when the mean amplitude of the beta power within the movement period (0–1.5 s relative to movement onset) was calculated and compared across stimulation conditions, no significant differences in beta ERD were observed (χ²(2) = 2.01, *p* = 0.365; Extended Data Fig. 2).

Taken together, these results indicate that behaviourally-driven closed-loop beta tACS modulates post-movement beta ERS in a temporally specific manner, enhancing beta power mainly within the stimulation-aligned window.

### Beta tACS increases retention of a visuomotor adaptation task

Having established that behaviourally-driven closed-loop tACS leads to an increase in beta ERS, we next wished to investigate whether this increase in activity drove changes in behaviour. As hypothesised, when the force field perturbation was applied, participants’ movement trajectories exhibited a systematic deviation from the straight line between the start point and the target (Fig. 1b, Fig. 4). An increase in lateral deviation was observed immediately upon transitioning to a different force magnitude and gradually decreased over subsequent trials within the same magnitude of applied force (Fig. 4).

**Figure 4.**
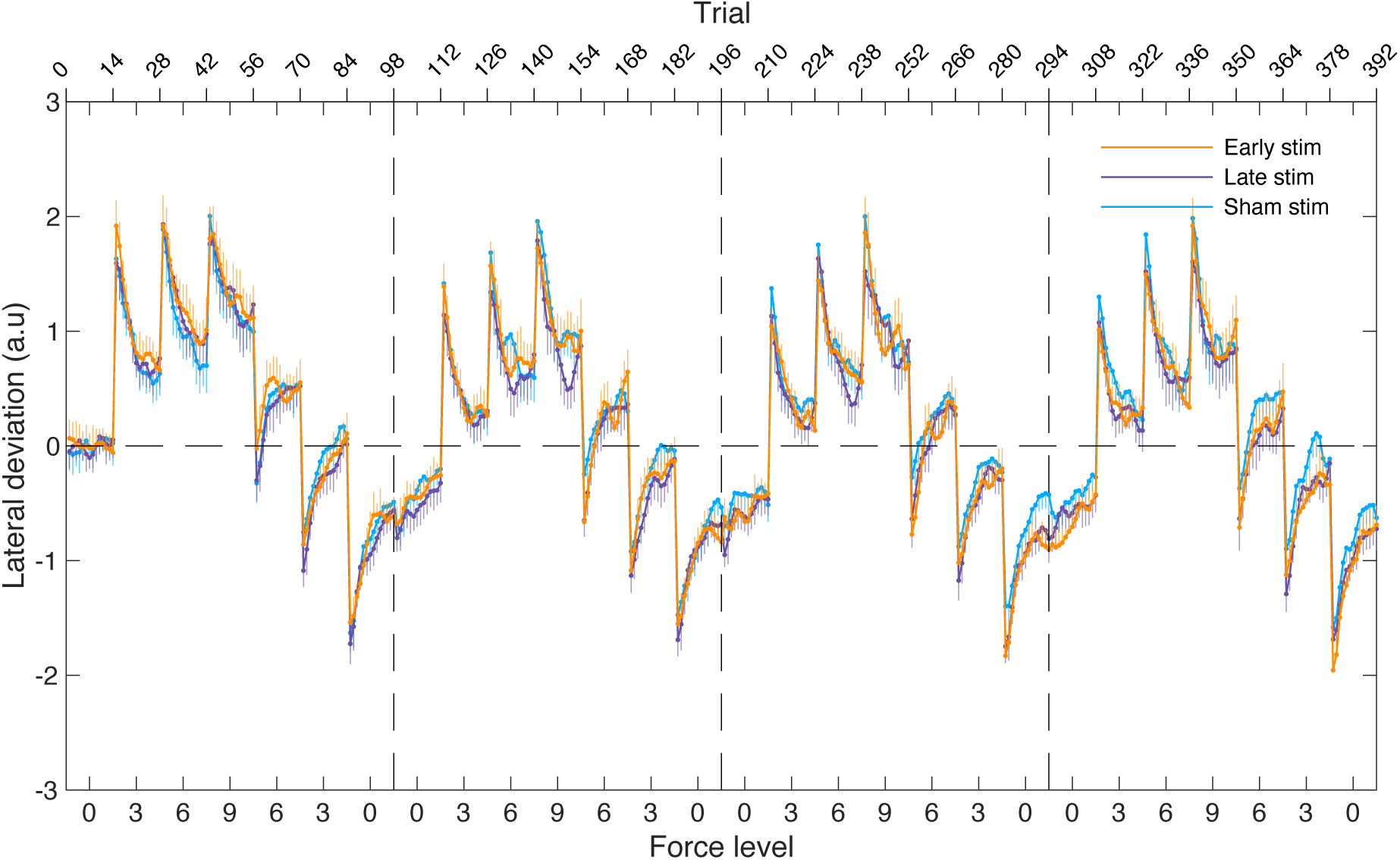
Trial-by-trial motor performance across stimulation conditions. Mean lateral deviation (± s.e.m) across all participants during the adaptation phase is shown across trials, as the force field changed in the sequence: 0-3-6-9-6-3-0, repeated for four blocks. Each force magnitude was applied for 14 consecutive trials. Positive values reflect deviations in the direction of the applied force (e.g., leftward deviation in response to a right-to-left force field). Negative values reflect deviations opposite to the force direction. Deviations were baseline-corrected using the mean from the initial zero-force trials of the first block.

Improvements on visuomotor adaptation tasks rely on two distinct behavioural processes: an adaptation to the new perturbation, which is thought to primarily be encoded in the cerebellum, and the retention of that new visuomotor transform, which is dependent on cortical regions ^22^. We therefore fitted a state-of-the-art, two-state model to capture the multiple processes ^23^. The model fitted well across all stimulation conditions (early: RMSE = 0.88 ± 0.03; late: RMSE = 0.86 ± 0.03; sham: RMSE = 0.86 ± 0.02; Fig. 5A). The estimated parameters revealed a significant effect of stimulation condition on the retention rate of the slow process (Aₛ, χ²(2) = 11.73, *p* = 0.003). Aₛ was significantly higher during early tACS compared to sham (*t*(24) = 2.850, *p* = 0.009, FDR-corrected), reflecting a tACS-induced enhanced retention of the slow process. Aₛ was also higher during late tACS compared to sham (*t*(24) = 2.955, *p* = 0.009, FDR-corrected). No difference was found between early and late tACS (*t*(24) = –0.105, *p* = 0.917, FDR-corrected; Fig. 5B). No significant differences between stimulation conditions were found for other model parameters (*p* values > 0.05 for A_f_, B_f_, and Bₛ; Fig. 5B), indicating that the effect of stimulation was specific to slow retention.

**Figure 5.**
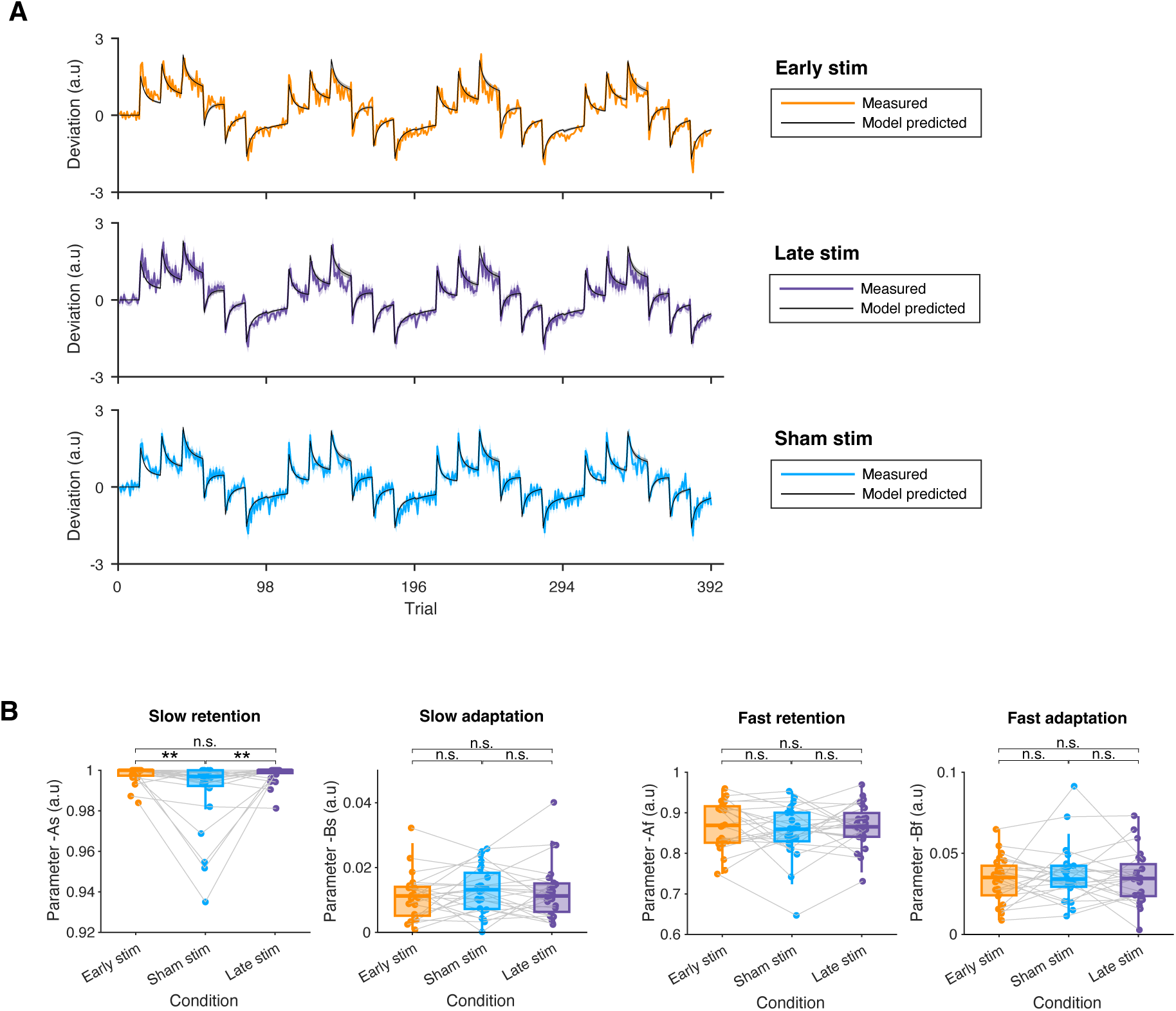
Two-state model fitting of motor adaptation across stimulation conditions. **(A)**. Measured (coloured) and two-state model-simulated (black) trial-by-trial deviations (mean ± s.e.m) during the motor adaptation task for early (orange), late (purple), and sham (blue) stimulation conditions. **(B).** Estimated model parameters across conditions: slow retention factor (*A_s_*), slow learning rate (*B_s_*), fast retention factor (*A_f_*), fast learning rate (*B_f_*). Box plots show the median, interquartile range, and individual data; grey lines link participant data across conditions. The slow retention parameter (*A_s_*) was significantly increased in both early and late stimulation conditions compared to sham, indicating enhanced retention. No significant differences were observed in the other parameters. n.s. non-significant, ** *p* < 0.01.

**Figure 6.**
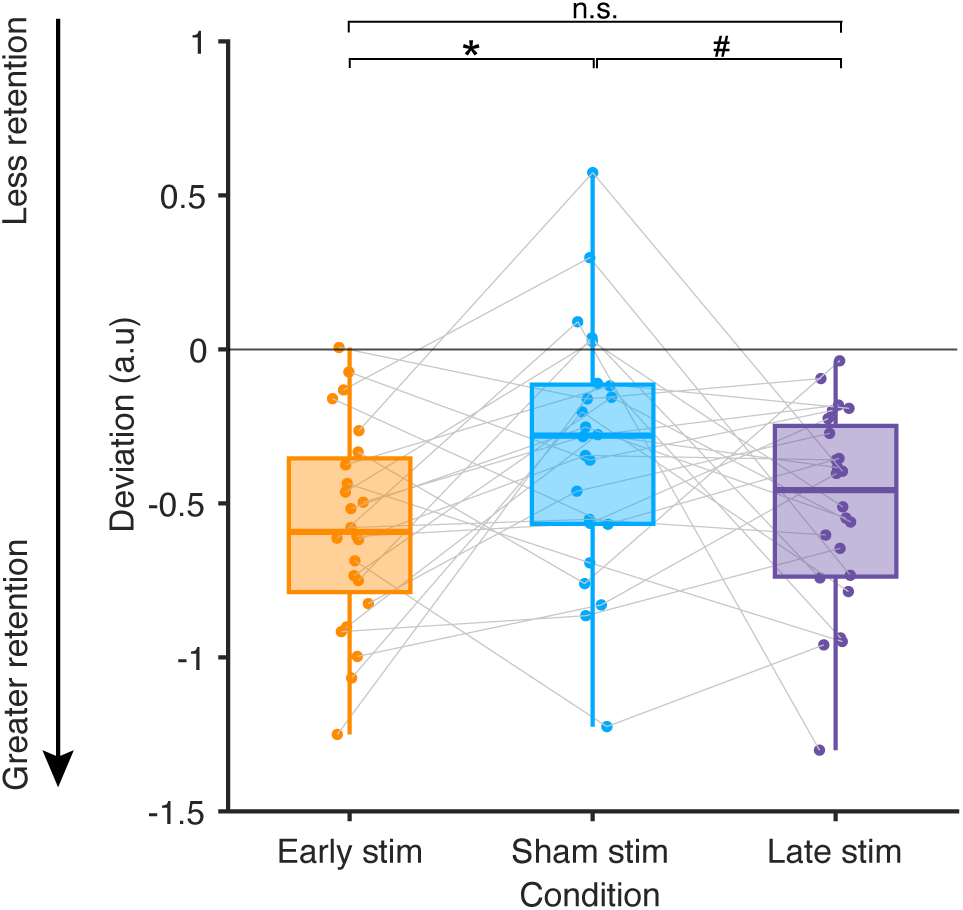
Measured retention across stimulation conditions. Motor performance measured as lateral deviation during the retention phase, shown for early stimulation (orange), sham (blue), and late stimulation (purple) conditions. More negative values indicate greater retention of the adapted state. Box plots show the median, interquartile range, and individual data; grey lines link participant data across conditions. Early stimulation resulted in significantly greater retention compared to sham and a marginal difference was also observed between late and sham stimulation. ^#^ *p* < 0.1, * *p* <0.05.

To confirm these findings from our model fit we also directly quantified retention in no-vision retention trials. Similarly to the model fit results, a significant effect of stimulation condition was observed (χ²(2) = 11.30, *p* = 0.003). During early tACS participants showed more negative lateral deviation, compared to the sham condition (*t*(24) = –2.772, *p* = 0.022, FDR-corrected), reflecting stronger retention of the previously adapted motor output. A marginal difference was observed between late stimulation and sham (*t*(24) = –2.157, *p* = 0.052, FDR-corrected). No significant differences were observed between early and late stimulation (*t*(24) = –0.615, *p* = 0.541, FDR-corrected). These results are consistent with the model-derived estimates, suggesting that closed-loop beta tACS enhanced both model-predicted retention and directly quantified retention.

### Enhanced beta ERS is associated with greater retention

We have shown that tACS leads to an increase in beta ERS amplitude, and to a specific improvement in retention. Finally, we wanted to determine whether these neural and behavioural effects were related. To address this, we included beta ERS power, computed across the broad post-movement window spanning both the early and late stimulation periods, as a predictor of retention in a linear mixed-effects model. Higher beta ERS was significantly associated with greater Aₛ values (β = 0.007, *p* = 0.045), suggesting that enhanced post-movement beta ERS correlates with increased retention of the novel visuomotor transform.

### Blinding of stimulation conditions was effective

Lastly, we wanted to ensure that neither participants nor the experimenter were able to distinguish between the three stimulation conditions. There was no evidence that reporting was better than chance (Participants: *p* = 0.266; experimenter: *p* = 0.205), with 16 out of 24 participants reporting the sham condition as active and the experimenter frequently reporting “don’t know” (Extended Data Figs. 3A-B), indicating successful participant blinding. No serious adverse events were reported. The most frequently reported sensations were mild tingling, itching, and sleepiness. There were no significant differences in the frequency of reported sensations across stimulation conditions (all *p* > 0.05, Extended Data Fig. 1C).

## Discussion

Here we have developed and implemented a novel behaviourally-triggered closed-loop tACS approach, which is capable of specifically increasing the post-movement-related beta ERS, with an associated increase in retention, compared with sham stimulation.

### Behaviourally-triggered closed-loop beta tACS increases movement-related beta ERS

We have shown that our behaviourally-triggered beta tACS increases movement-related beta ERS. Importantly, our EEG analyses only included trials without concurrent stimulation, ensuring that the observed increases in beta power reflect changes in brain activity without being contaminated by stimulation artefacts. While tACS commonly induces entrainment during active stimulation (“online” effects) and can outlast the stimulation, the temporally specific, offline enhancement of beta ERS observed here is more consistent with synaptic plasticity mechanisms, such as spike-timing-dependent plasticity ^24,25^. This interpretation is supported by previous work using intermittent, brief alpha tACS, which found similar after-effects regardless of whether stimulation was phase-continuous or phase-discontinuous ^26^. These findings suggest that the offline effects we observe here may be driven by synaptic plasticity rather than mere entrainment.

Previous work has demonstrated offline effects of continuous beta tACS but the findings remain inconsistent ^27–30^. Here, we show that even brief (2 seconds) trains of beta tACS can induce specific offline neural effects. This finding contrasts with studies using alpha tACS, where longer trains (8 seconds) but not shorter trains (3 seconds) induced offline effects ^26^. We believe that this inconsistency is likely because the effectiveness of brief stimulation critically depends on the specific brain or behavioural state at the moment of stimulation ^31,32^. For instance, even a 1 second tACS train during sleep can significantly modulate spindle activity ^32^. Similarly, our closed-loop stimulation delivered immediately after movement offset, when the brain naturally exhibits increased beta activity, likely preferentially enhances cortico-subcortical motor plasticity and reinforces functionally relevant motor network activity.

Behaviourally-triggered closed-loop beta tACS specifically improves cortically-dependent behaviour performance on a visuomotor adaptation task. This task relies on two distinct but concurrent behavioural processes, requiring the participant both to adapt to the new perturbation, which is primarily encoded in the cerebellum, and retain that new visuomotor transform, a cortically-dependent process ^33–35^. In line with previous brain stimulation studies we therefore hypothesised that cortical tACS applied above M1 would lead to a specific effect on retention while having no effect on adaptation ^33,36^. Our behavioural results support this hypothesis. We observed that beta tACS applied over M1 shortly after movement offset significantly improved retention. These findings align with existing evidence emphasizing the crucial role of M1 in motor retention. For example, studies using transcranial magnetic stimulation (TMS) have shown that reducing M1 excitability shortly after adaptation disrupts retention, suggesting M1 is vital for early memory consolidation ^37,38^. Conversely, studies indicate that enhancing M1 excitability utilizing anodal transcranial direct current stimulation (tDCS) immediately after adaptation improves retention ^33,36^. Our findings extend these studies by demonstrating that short, behaviourally-triggered bursts of beta tACS delivered to M1 can also enhance retention. Due the short, but neurophysiologically-relevant, nature of the stimulation, the same total duration of stimulation can be applied over more movement repetitions, compared to conventional continuous stimulation.

Indeed, previous work highlights a temporally sensitive window immediately after training, during which the motor cortex is particularly receptive to neuromodulation. For instance, single-pulse TMS applied immediately after trial completion disrupts retention, whereas stimulation applied slightly later (700 ms post-trial) does not have the same disruptive effect ^38^. Our data suggest that while immediate post-movement stimulation (i.e., early stimulation) strongly enhances retention, stimulation applied at a later period (i.e., late stimulation) also has a marginal benefit on behaviour. A plausible explanation for this could be that our task led to a relatively prolonged post-movement beta ERS compared with hand-held joystick tasks. Thus, even stimulation applied slightly later may interact beneficially with ongoing neural processes related to motor retention.

However, we only assessed retention during a single session and did not examine longer-term behavioural effects (e.g., 24-hour afterwards). Therefore, it remains to be tested whether the observed improvements in retention would persist over time and whether repeated stimulation sessions could enhance or sustain the benefits.

### Movement-related beta ERS amplitude is correlated with cortically-dependent retention

In our study, applying beta tACS immediately after the movement increased post-movement beta ERS, and this increase was associated with larger retention factor in the slow process. Our results align with recent findings demonstrating that 20 Hz tACS over the motor cortex after training enhances motor performance measured after 24-hours and 7-days compared to sham stimulation^39^. Similarly, previous studies have shown that beta-tACS increases corticomuscular coherence, which positively correlates with improved motor skill retention ^18^. The relationship between beta ERS and retention was also observed in observational studies. For example, healthy individuals who exhibit larger increases in beta ERS during practice typically show stronger retention of motor skills the next day. In contrast, patients with Parkinson’s disease show smaller beta ERS and impaired motor skill retention ^40^.

Mechanistically, a stronger beta ERS may indicate a brain state that supports memory retention by reinforcing the current motor plan and reducing the need for further corrections. Previous work has proposed that beta activity increases when the brain is trying to maintain a stable motor state^41^. During learning, a strong beta ERS after movement indicates more confidence in the motor output or the internal forward model. Conversely, a weak beta rebound might represent a need for further updating or correction ^6^. Therefore, stronger beta ERS following beta-tACS was associated with greater retention of motor output.

### Towards behaviourally-triggered closed-loop stimulation for real-world settings

We have demonstrated the ability of closed-loop stimulation to specifically interact with key neural dynamics, resulting in behavioural improvement. This has many potential applications as a putative therapeutic tool, but to be successful, it needs to be practicable to achieve. Conventional closed-loop brain stimulation approaches typically rely on continuous EEG monitoring, real-time neural decoding, and complex artefact removal ^7,19,42–44^, limiting their utility for clinical and practical applications, particularly in noisy or mobile settings. Our approach simplifies these challenges by using wearable motion sensors and mobile EEG, designed to directly translate into the home setting if required. Triggering the tACS via behaviour means that we can accurately target behaviourally-relevant neural activity and avoid the need for continuous EEG monitoring and artefact correction.

This approach is also highly efficient with a total stimulation duration per session of ∼8.7 minutes. This duration is substantially less than that of conventional tACS protocols lasting 20–30 minutes ^45–49^, but is still capable of robustly enhancing post-movement beta ERS, thereby reducing the potential risks and burden to the participant.

## Conclusions

Here, we designed and implemented a portable, closed-loop neuromodulation system using motion sensor-based triggering of beta-frequency tACS. We demonstrated that behaviourally-triggered closed-loop tACS can effectively modulate specific aspects of neural activity and improve retention on a visuomotor adaptation task. This technique therefore opens a novel putative therapeutic route to improve motor function in clinical populations characterised by impaired movement-related beta activity, such as people with Parkinson’s disease or stroke.

## Methods

### Participants

Twenty-seven healthy participants (18-35 years old, 17 females) were recruited for this study. All participants were right-handed according to the Edinburgh Handedness Inventory and had normal or corrected-to-normal vison ^50^. All participants fulfilled the following inclusion criteria: (1) no history or current neurological or psychiatric disease; (2) no impairment of upper limb function; and (3) no use of drugs affecting the central nervous system. Safety screening questionnaires for brain stimulation were completed prior to the experiment. Written informed consent was obtained from all participants in accordance with Declaration of Helsinki. This study was approved by the Central University Research Ethics Committee, University of Oxford (Ethics Reference: R81071/RE003). Three participants withdrew before completing all experimental sessions (one reported scalp skin intolerance following the second experimental session and two dropped out due to personal reasons). Twenty-four participants completed the experiment and were included in the analyses.

### Study design

This within-subject, double-blinded, sham-controlled study included three sessions, with the only difference between sessions being the stimulation condition. The order of stimulation was counterbalanced across participants using a randomization code. Both participants and the experimenter, interacting with the participants, were blinded to the stimulation allocation. A second experimenter, responsible for technical implementation, was not involved in participant interaction and did not disclose stimulation details. Sessions were separated by at least seven days (median 7 days; interquartile range [IQR] 7–9.5).

### Motor adaptation task and behavioural data acquisition using Kinarm

In each session, participants performed a target-reaching task with their right hand using a Kinarm END-Point robotic system (Kinarm, Kingston, Canada). Participants were seated in a height-adjustable chair in front of the Kinarm and faced a horizontal mirror that reflected visual stimuli from a downward-facing monitor. The right Kinarm handle was displayed on the screen as a white circle (radius: 0.5 cm). Handle position (x, y) and velocity in the horizontal plane were recorded at a sampling rate of 1000 Hz using the Kinarm system.

The task required participants to use the right Kinarm handle to make straight, planar reaching movements between two target points spaced 15 cm apart, alternating between outward (away from the body) and inward (toward the body) directions. At the start of each trial, participants positioned the handle at the start point. Once the handle was held at this position, the target turned yellow, indicating a go cue. Participants were instructed to move as quickly and accurately as possible to the target. Upon reaching it, the target turned purple and then became the start point for the next trial. This ensured an alternation of inward and outward movements. An inter-trial interval of 5 seconds was used to quantify the post-movement beta ERS.

The movement task in each session consisted of three sequential phases: practice, adaptation, and retention. During practice, participants completed 14 trials without any force field perturbation, and feedback was given if the movement was performed not fast enough. The adaptation phase included a total of 392 trials, divided into four blocks of 98 trials each. A velocity-dependent lateral force field was introduced during this phase, applying force along the x-axis (perpendicular to movement direction). The magnitude of the force field changed every 14 trials following this sequence: *0, 3f, 6f, 9f, 6f, 3f, 0* (Fig. 1A), see section ‘Two-state model fitting’ for more details. Participants were not informed about the presence of the force perturbations. The direction of the force perturbation (leftward or rightward) remained constant within a session but varied across sessions. The order of the force direction across three sessions (i.e., left-right-left or right-left-right) was counterbalanced across participants and stimulation condition. Both during practice and adaptation phases, continuous visual feedback of the handle position was provided. Immediately after the adaptation phase, a retention phase of 14 trials was conducted. In these trials, no force was applied and the visual feedback of both the handle was removed during movement. This phase allowed assessment of the retention of the adapted motor response in the absence of visual feedback.

### Closed-loop transcranial alternating current stimulation (tACS)

Kinematic data were also acquired using a single VIVE Tracker 3.0 and two HTC VIVE Base Stations (HTC, Taiwan). Kinematic data from the VIVE Tracker were used to trigger the brain stimulation. We opted to trigger brain stimulation from the kinematic data obtained from the VIVE Tracker rather than the Kinarm, as the VIVE Tracker is more affordable and portable, allowing for future studies outside the laboratory. The VIVE Tracker rigidly attached to the Kinarm handle and remained in place throughout the entire study and coordinate systems of the Kinarm robot and the VIVE Tracker were aligned. VIVE Tracker positional data were wirelessly streamed via a VIVE Dongle at a sampling rate of 90 Hz. A custom C++ application (ViveTrackQt) continuously recorded the VIVE tracker’s position and interfaced in real time with a MATLAB script via a TCP/IP connection. When the VIVE tracker (attached to the Kinarm handle) had entered the target area (defined as time *t₀*) and remained continuously within this area for at least 150 ms (defined as time *t₁,* Fig. 1C) stimulation was triggered.

TACS was applied during the adaptation phase but not the practice or retention phases. tACS was delivered using a Starstim 8 system (Neuroelectrics, Barcelona, Spain). Two 1π cm² Ag/AgCl electrodes (Starstim NG Pistim) were integrated into an EEG cap, with the stimulation electrode placed over the left motor cortex (C3) and the return electrode positioned at Pz according to the standard 10–20 system. Stimulation was applied at 20 Hz, with the current amplitude of ±2 mA peak-to-peak. Electrode impedance was maintained below 10 kΩ throughout each session.

Three distinct stimulation conditions were used. In the early stimulation condition, 20 Hz tACS was delivered immediately following this 150 ms hold period (*t₁*) and lasted for 2 seconds. In the late stimulation condition, stimulation was delivered 2.5 seconds after *t₁* and also lasted 2 seconds. In the sham condition, stimulation was similarly triggered 2.5 seconds after t₁ but lasted only 100 ms. All stimulation trains included 50 ms ramp-up and ramp-down periods. tACS was applied in four out of every six trials, with the remaining two trials left stimulation, and thus artifact-free for EEG data analysis.

### Behavioural data analysis

The velocity signals recorded from the Kinarm system were low-pass filtered using a fourth-order Butterworth filter with a 30 Hz cutoff. Movement onset was defined as the time point when velocity exceeded a threshold, calculated as the mean plus three standard deviations (SD) of the velocity during the “rest” period and remained above this threshold for at least 100 ms ^51,52^. Similarly, movement offset was defined as the time before the velocity fell below this threshold and remained there for at least 100 ms. Onset and offset were identified within each trial. The interval from movement onset to movement offset was defined as the movement period for each trial.

Movement performance was quantified by the maximum lateral deviation, defined as the peak perpendicular distance between the Kinarm handle trajectory and the straight line connecting the start point and target. This was achieved by using the *findpeaks* function in MATLAB, with a minimum peak width of 100 ms. Deviation sign was defined relative to the direction of the applied force field direction: deviations toward the force direction were considered positive, while deviations opposite the force direction were negative. Although no force was applied during the retention phase, the deviation sign was defined relative to the force direction used in the previous adaptation phase.

To account for individual baseline biases, lateral deviation values were baseline-corrected by subtracting the mean deviation during the initial zero-force trials of the first block (Fig. 1A). Measured retention was defined as the mean lateral deviation across all trials in the retention phase.

### Two-state model fitting

To further characterize motor adaptation and retention processes, we fitted a two-state model to the trial-by-trial lateral deviation data in the adaptation phase ^23^. In this model, the motor output 𝛸(𝑛) on trial 𝑛, is the sum of a fast process 𝛸_!_(𝑛) and a slow process 𝛸_*f*_(𝑛). These two processes update independently based on the error 𝑒(𝑛 − 1) and motor output 𝛸(𝑛 − 1) on the previous trial following:

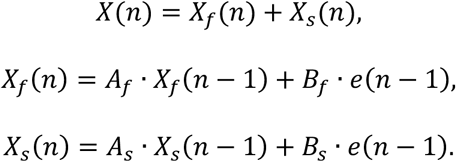

Here, 𝐴_*f*_ and 𝐴_*s*_ are retention factors, and 𝐵_*f*_ and 𝐵_*s*_ are learning (adaptation) rates for the fast and slow processes, respectively. All parameters were constrained to the interval [0, 1], with the additional constrains 𝐴_*f*_ < 𝐴_*s*_ and 𝐵_*f*_ > 𝐵_*s*_, reflecting the faster learning and lower retention of the fast process compared to the slow process.

The error 𝑒(𝑛), i.e., lateral deviation, was defined as the difference between the scaled force field and the motor output on each trial:

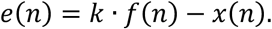

where 𝑓(𝑛) is the applied force magnitude on trial 𝑛 and 𝑘 is a free scaling parameter that maps the force magnitude to the measured lateral deviation. Force magnitudes were encoded as 0 (no force), 1 (force magnitude 9), and linearly interpolated for intermediate values (e.g., 1/3 for force magnitude 3 and 2/3 for magnitude 6).

Model fitting was performed by simulating the entire sequence of trials and minimizing the sum of squared errors between predicted and observed deviations using MATLAB’s *fminsearch* function. Firstly, a group-level fit was conducted by concatenating all trials across participants and sessions, allowing estimation of five free parameters: 𝐴_*f*_, 𝐵_*f*_, 𝐴_*s*_, 𝐵_*s*_, and the scaling factor 𝑘.

Secondly, individual-level fits were performed separately for each participant and session. For these individual fits, the four learning parameters (𝐴_*f*_, 𝐵_*f*_, 𝐴_*s*_, 𝐵_*s*_) were initialized using the group-level estimates, while the scaling factor 𝑘 was held fixed to the group-level value. This procedure yielded participant-and session-specific estimates of the learning and retention. The goodness-of-fit was assessed using the root mean square error (RMSE).

### EEG recording and analysis

EEG data were continuously recorded during the movement task using a 64-channel wireless system (Smarting®, mBrainTrain®, Belgrade, Serbia) with an elastic EEG cap (EASYCAP, Herrsching, Germany). Electrodes were positioned according to a standard 10-20 system, with FPz as ground electrode and FCz as the reference electrode. Electrode impedances were maintained below 10 kΩ throughout each session. EEG signals were digitized at a sampling rate of 250 Hz.

Data preprocessing was performed using EEGLAB (version 2024.0) ^53^ and MATLAB (R2023a). Raw data were band-pass filtered between 1 Hz and 45 Hz using a Butterworth Infinite Impulse Response (IIR) filter (zero phase shift, filter order: 6). The data were epoched from −2 to +7 seconds relative to movement onset. Epochs containing stimulation artefacts were rejected. Then, bad channels with a leptokurtic voltage distribution were replaced by interpolating between the surrounding electrodes (spherical spline interpolation; mean ± SD: 3 ± 3 channels). Next, independent component analysis (ICA) with the Second Order Blind Identification (SOBI) algorithm was performed, and artefactual components were identified and discarded (mean ± SD: 13 ± 6 components) using the EEGLAB plugin ICLables ^54^.

Following artefact rejection, data were re-epoched for subsequent analyses: from –1.5 to +3 seconds relative to movement onset (for beta ERD) and from –1.5 to +5 seconds relative to target reached (for beta ERS). Time–frequency analysis was performed using Morlet wavelets in FieldTrip (version 20240614). Spectral power was estimated from 5 to 40 Hz (1 Hz steps), with a temporal resolution of 20 ms, using a fixed wavelet width of 9 cycles. Power was computed on a single-trial basis. Baseline correction was applied from –0.5 to –0.2 seconds relative to movement onset, and the results were converted to decibel (dB) units to normalize power.

Beta ERS was quantified within stimulation-aligned time windows: an “early stimulation” window from 0 to 2 seconds and a “late stimulation” window from 2.5 to 4.5 seconds relative to the time of movement offset (*t₁*, see ‘Closed-loop tACS’). Beta ERD was computed from 0 to 1.5 seconds relative to movement onset. Analyses focused on 15-30 Hz and were restricted to electrodes over the contralateral sensorimotor cortex, including C1, C5, CP1, CP3, CP5, P1, P3, and P5, located between the two stimulation electrodes (C3 and Pz).

### Statistical Analyses

Clusters showing beta power difference between stimulation conditions were identified using non-parametric cluster-based permutation tests (1000 permutations), with correction for multiple comparisons based on the temporal cluster size criterion ^55^.

To assess the stimulation condition effects on behavioural and EEG outcomes, linear mixed-effects models (LMMs) were implemented using the lme4 package in R. Separate models were constructed for beta ERS power averaged from 0 to 4.5 seconds following movement offset, covering both the early and late stimulation time windows, beta ERD power, measured retention (lateral deviation during the retention phase), or model-derived parameters. Each model included stimulation condition (early, late, sham) as a fixed effect and participant as a random intercept. When a significant main effect was observed, post hoc pairwise comparisons were performed with false discovery rate (FDR) correction. Additional LMMs were used to test whether model parameters could be predicted by beta ERS.

The effectiveness of participant and experimenter blinding was assessed using Fisher’s exact test to determine whether the distribution of perceived stimulation types differed across stimulation conditions. Similarly, sensations to stimulation (e.g., itching, tingling) were summarized descriptively and condition differences were also evaluated using Fisher’s exact test.

## Author contributions

MW: conceptualisation, data acquisition, methodology, data analysis, writing – original draft, writing – review and editing. ZX: conceptualisation, data acquisition, writing – review and editing. MKF: conceptualisation, data acquisition, supervision, writing – review and editing. NS: methodology, writing – review and editing. LB: data acquisition, writing – review and editing. FT: data acquisition, writing – review and editing. PLW: data acquisition, writing – review and editing. CN: experimental design, formal analysis, writing – review and editing. AS: experimental design, writing – review and editing. CZ: conceptualisation, experimental design, methodology, formal analysis, supervision, funding acquisition, and writing review and editing. CJS: conceptualisation, methodology, resources, funding acquisition, supervision, and writing – review and editing.

## Data availability

The datasets and code will be shared via the MRC BNDU data sharing platform on ^40^.

## Rights retention text

This research was funded by the Wellcome Trust [203139/Z/16/Z]. For the purpose of open access, the author has applied a CC BY public copyright licence to any Author Accepted Manuscript version arising from this submission.

## Extended Data

**Extended Data Figure 1.**
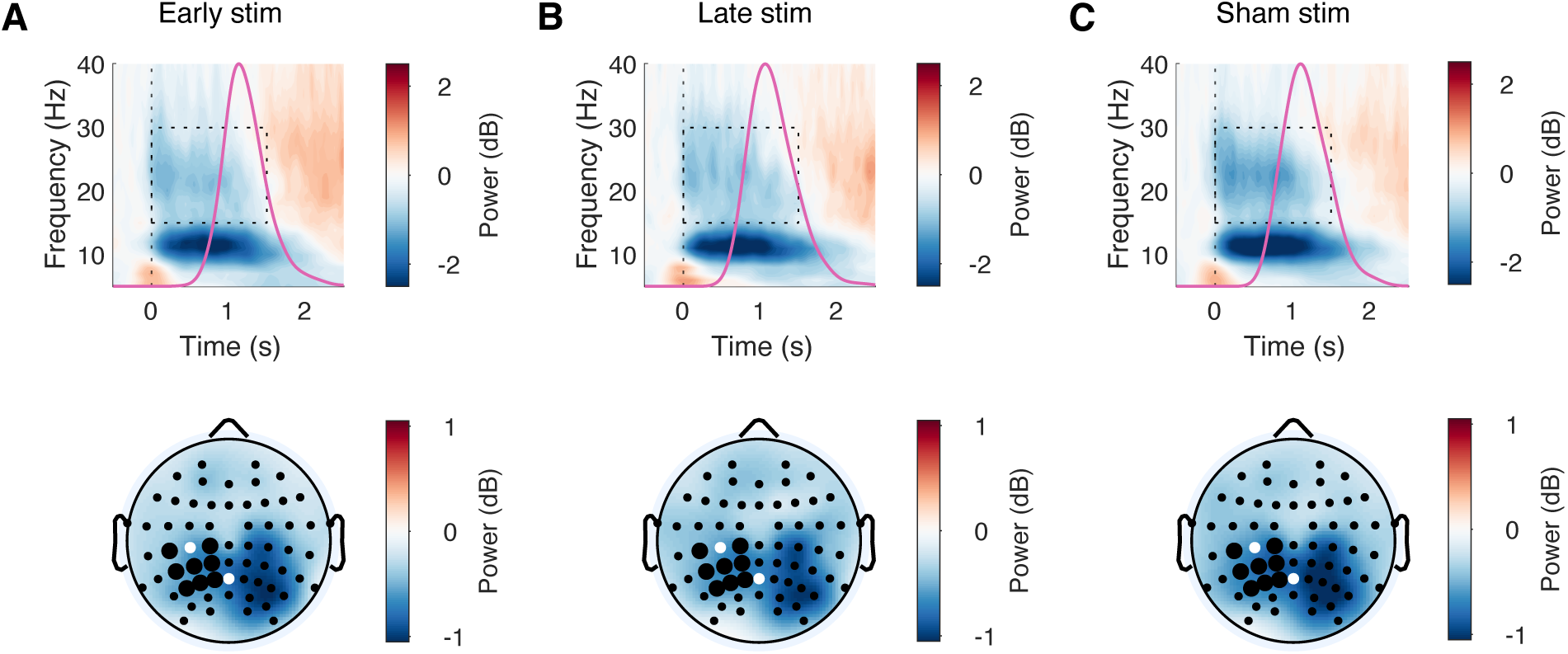
Time–frequency responses during movement across stimulation conditions. **(A)**. Time-frequency response aligned to movement onset (*t_0_ = 0 s*) for the early stimulation condition. The power was averaged across sensors covering the contralateral sensorimotor area (highlighted in bold in bottom panel). The magenta curve shows the distribution of movement time across participants. The black rectangle indicates the time–frequency window used to quantify beta ERD (0–1.5 s, 15–30 Hz). The scalp topography shows bilateral distribution of beta ERD. Black dots mark electrodes in the predefined sensorimotor area; white dots indicate stimulation electrodes (C3 and Pz). **(B).** Similar as (A), for the late stimulation condition. **(C).** Similar as (A), for the sham stimulation condition.

**Extended Data Figure 2.**
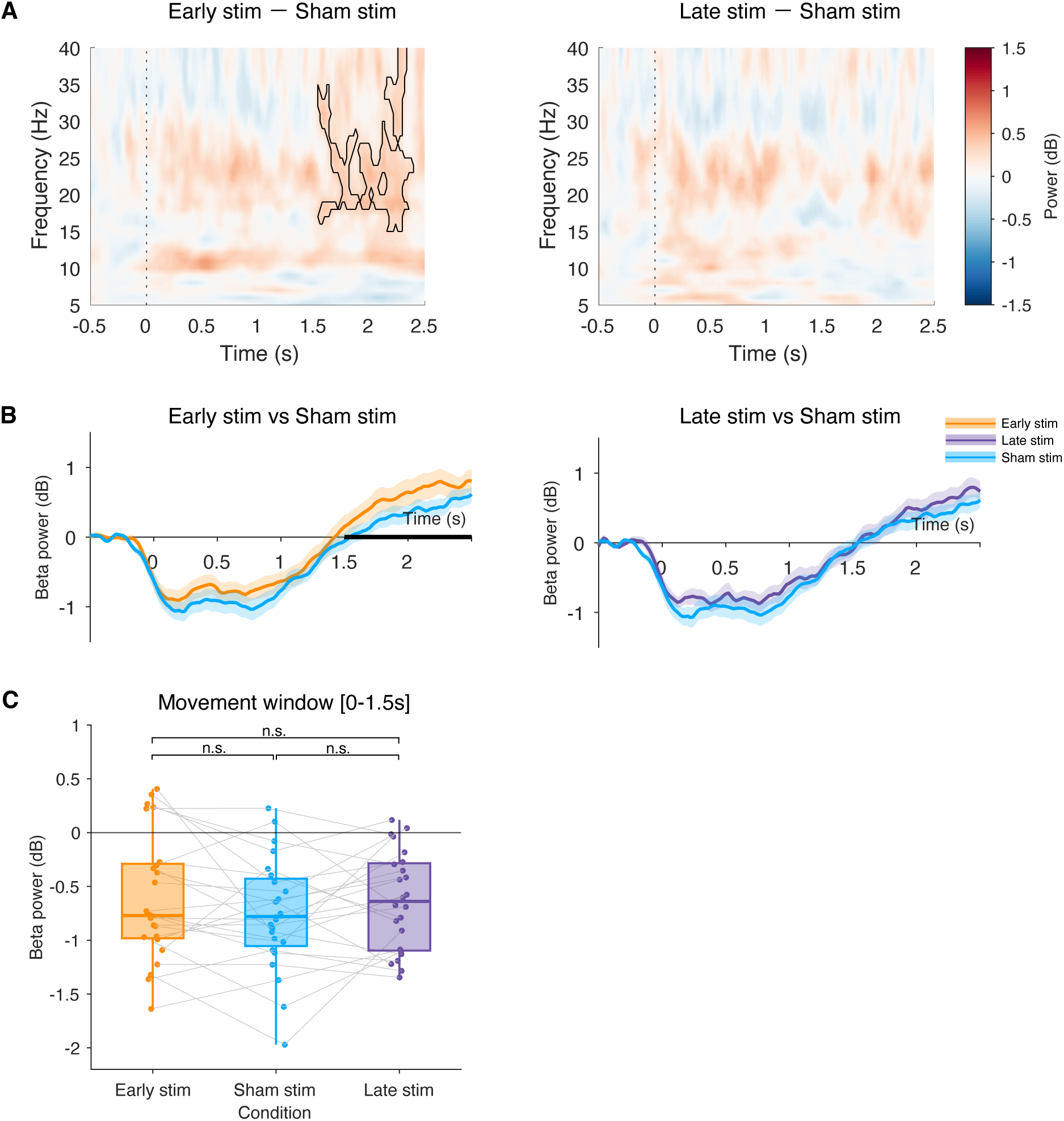
Modulation of beta activity during movement by stimulation conditions. **(A)**. Time frequency response difference in the sensorimotor cortex between conditions. The black outlines indicate significant clusters with *p* < 0.05 by cluster-based permutation test. **(B).** Time course of beta power (15–30 Hz) averaged over the contralateral sensorimotor area, aligned to movement onset (*t₀* = 0 s). Left: comparison between early (orange) and sham stimulation (blue). Right: comparison between late (purple) and sham (blue) stimulation. Black horizontal bars indicate time intervals with significant differences between conditions, determined by cluster-based permutation testing (*p* < 0.05). **(C).** Mean beta power extracted from movement onset to 1.5 s afterwards. Each dot represents an individual participant; grey lines link participant data across conditions. Box plots show the median and IQR. No significant difference on beta ERD was observed across three stimulation conditions.

**Extended Data Figure 3.**
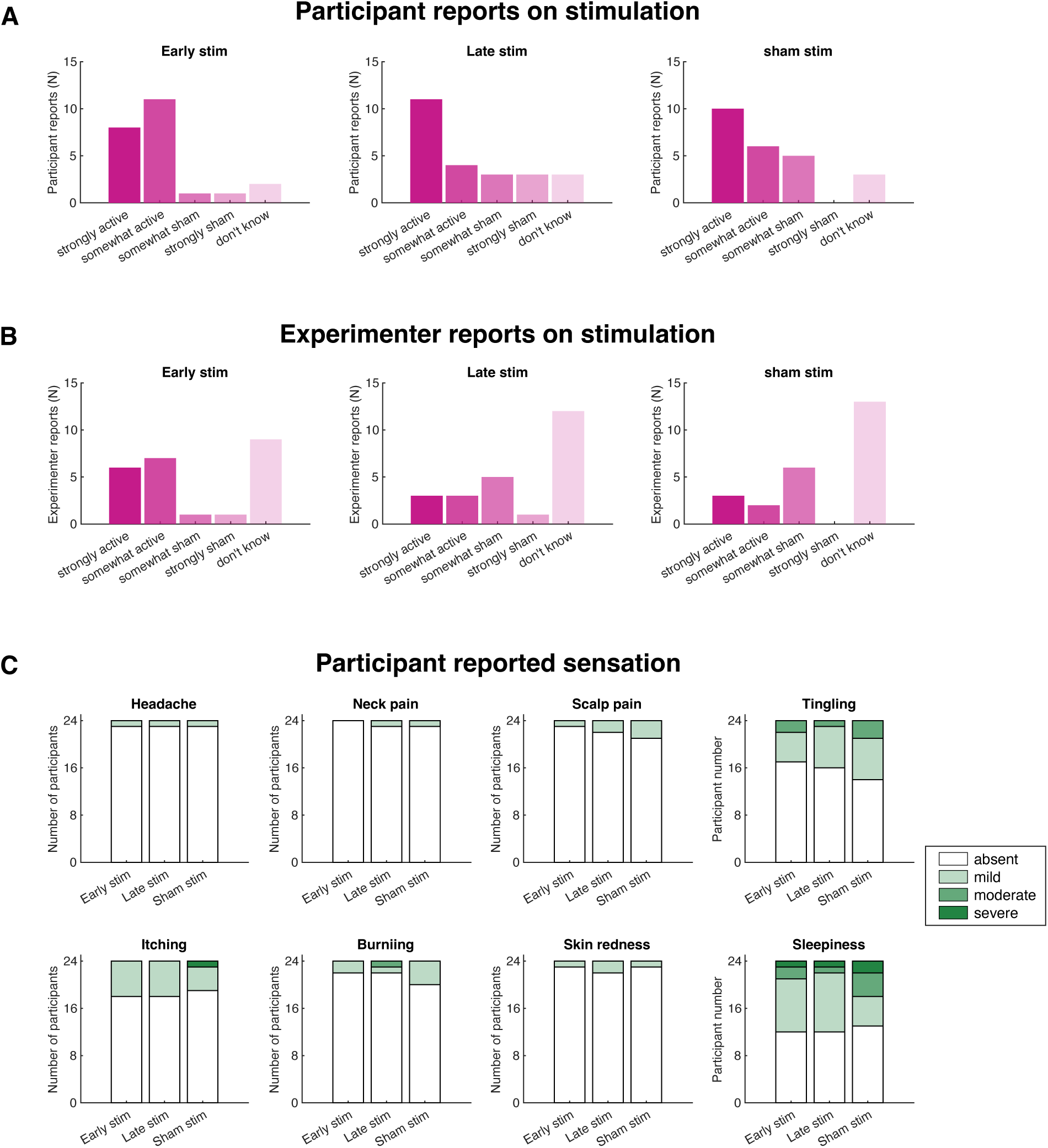
Blinding effectiveness and reported sensations across stimulation conditions. **(A-B).** Distribution of reported stimulation type from participants and experimenter in early, late, and sham stimulation conditions. Bars show the number of participants/experimenter reporting the stimulation as strongly active, somewhat active, somewhat sham, strongly sham, or don’t know. **(C).** Participants’ reported sensations during stimulation. Each panel corresponds to one sensation type (e.g., tingling, itching, sleepiness). Stacked bars indicate the number of participants reporting each sensation at four severity levels: absent, mild, moderate, and severe.

